# *Candida auris* Metabolism and Growth Preferences in Physiologically Relevant Skin-like Conditions

**DOI:** 10.1101/2025.05.18.654780

**Authors:** Jonathan P. Nicklas, Clay Deming, ShihQueen Lee-Lin, Sean Conlan, Zeyang Shen, Ryan A. Blaustein, Julia A. Segre

## Abstract

*Candida auris* is an opportunistic, multidrug-resistant yeast with high capacity of skin colonization in healthcare settings, which can lead to subsequent infections with high mortality rates. Given the recent emergence of at least four distinct clades at the global scale, little remains known about how *C. auris* is so adept at growing on skin and the key genes and pathways it utilizes to metabolize the scarce nutrients available. Here, we identify the roles that conventional and alternative carbon metabolism genes and metabolic pathways have in facilitating *C. auris* growth through laboratory-based experiments and bioinformatics analyses. In artificial skin-like media, all four clades of *C. auris* were more capable of growing than *C. albicans* SC5314, a clinically relevant counterpart. By investigating the differential regulation of *C. auris* when growing in skin-like media as compared to rich fungal media, we uncovered hundreds of genes in multiple metabolic pathways. To further test the mechanisms of these metabolic pathways, we deleted several non-essential gene candidates including *FOX2* (B9J08_002847), *CAT2* (B9J08_000010), and *ICL1* (B9J08_003374). The mutant strains all exhibited abrogated growth in skin-like media and demonstrated nutrient preferences that differed from the wild type. Thus, we propose a model of how *C. auris* has the capacity to metabolize nutrients that are naturally available on skin by changing its metabolic profile. Targeting these metabolic pathways to mitigate *C. auris* growth on skin is a potential avenue to explore in controlling the spread of this emerging human fungal pathogen.

**IMPORTANCE:** *Candida auris* is an emerging fungal pathogen with skin as its primary site of colonization and subsequent transmission. Here, we show the importance of conventional and alternative carbon metabolism for *C. auris’* ability to grow in artificial skin-like media. This knowledge provides a better understanding of *C. auris* metabolism and sheds light on genes and pathways that could be targeted to interfere with persistent skin colonization.

## INTRODUCTION

*Candida auris* is a yeast that was first identified in a clinical report in a Japanese hospital in 2009 and has since independently emerged across four continents (1, 2). Epidemiological and genetic analysis categorized these four clades as: South Asia: Clade I, East Asia: Clade II, South Africa: Clade III, and South America: Clade IV (1). Tens to hundreds of single nucleotide polymorphisms (SNPs) may exist between two isolates of the same clade, but thousands of SNPs distinguish clades. New reports have identified a potential Clade V in Iran (3), and a potential Clade VI in Bangladesh (4) and Singapore (5). The origin of *C. auris* prior to colonizing humans and its recent emergence remains a topic of deep interest (6). *C. auris* has emerged as a cause of outbreaks in healthcare settings such as nursing homes and hospitals where it can persist for extended periods of time in rooms with colonized patients on high touch surfaces like mattresses, tables, chairs, floors, and doorknobs (7-10). *C. auris* exhibits resistance to disinfectants typically used in healthcare settings (11). *C. auris* can produce invasive candidiasis of the blood which can lead to life-threatening disease for individuals with various risk factors including compromised immunity, diabetes, stroke, indwelling medical devices, long term use of antibiotics or antifungals, and recent surgery (12, 13). The mortality rate for *C. auris* invasive infections is estimated to be 30-60% (12). A 2020 systematic review and meta-analysis evaluated over 4,700 cases reported in at least 33 countries and showed that *C. auris* (Clades I-IV) exhibit a high antifungal resistance profile of 91% resistance to fluconazole, 12% resistance to amphotericin B, 12.1% resistance to caspofungin, 0.8% resistance to micafungin, and 1.1% resistance to anidulafungin (13).Taken together, the challenges posed by *C. auris* contributed to its designation as an Urgent Threat in the CDC’s 2019 Antibiotic Resistance Threats Report in the US (14) and as a Critical Priority in the WHO’s 2022 Fungal Priority Pathogens List (15).

Skin is considered the primary reservoir for *C. auris* which differs from other *Candida* species that are more frequently associated with the gastrointestinal, oral, and urinary tracts (16). Vallabhaneni et al., 2017 evaluated the first *C. auris* infection cases in the USA reported to the CDC and they identified *C. auris* on human skin in the groin and axilla as well as the nares and rectum of patients (10). Eyre et al., 2018 found *C. auris* on the axilla and groin of patients in a neurosciences ICU at Oxford University Hospital, UK where an outbreak occurred from 2015-2017 (17). Adams et al., 2018 detected the presence of *C. auris* on the nares, axilla, and groin of patients in healthcare facilities in New York City, New York, USA in 2017 (7). Sexton et al., 2021 found *C. auris* in bilateral axillary and inguinal composite skin in a ventilator-capable unit in Chicago, Illinois, USA in 2018 (9). Proctor et al., 2021 screened ten body sites of residents of a skilled nursing facility in Chicago, Illinois over a three month point prevalence survey and found that *C. auris* prevalently colonized anterior nares, palm and/or fingertips, and toe web (18). Proctor et al. also discovered that residents could be colonized discreetly or simultaneously at multiple skin sites while validating *C. auris’* propensity for skin colonization and growth (18). Currently, CDC recommends that healthcare providers screen the axilla and groin to identify *C. auris* (https://www.cdc.gov/candida-auris/screening/index.html). Taken together, these studies show that *C. auris* persists on skin which predisposes a patient to infection (16). Moreover, long term skin colonization also favors transmission from patient to patient and overall spread in healthcare settings (19).

Several simulation studies and animal models have demonstrated that *C. auris* can grow and persist for extensive periods under skin-like conditions. Eix et al., 2022 demonstrated that *C. auris* could grow in a formulation of synthetic sweat medium for 24 hours and form a biofilm (20). Horton et al., 2020 demonstrated with a pig skin model supplemented with synthetic sweat medium that *C. auris* forms higher burden biofilms than *C. albicans* (21). Johnson et al., 2021 showed that *C. auris* could persist on pig skin for days at a time when grown in synthetic sweat media and that it can grow near the hair follicle (22). Huang et al., 2021 also demonstrated that *C. auris* could reside in murine skin within the hair follicle/sebaceous gland for months after the skin surface swabs tested negative (23). Guolei et al., 2025 showed that *C. auris* can coat the murine hair shaft and reside in the hair follicle (24). Santana et al., 2023 discovered an adhesin, *SCF1*, that is specific to *C. auris* that is critical for human and mouse skin colonization (25). Shivarathri et al., 2024 also showed that *Hog1* mitogen-activated protein kinase is essential for efficient skin colonization (26). Advancing understanding of the genes and metabolic pathways involved in *C. auris* fitness in these types of scenarios may help facilitate development of more effective strategies for control.

Skin is a nutrient poor environment that is not particularly hospitable to an incoming pathogen (27). Skin also has a range of distinct microenvironments: sebaceous, dry, moist, and foot (27). Each skin microenvironment has its own nutrient profile based on sweat and sebum gland densities which vary depending on body site (27). Sweat glands (eccrine and apocrine) secrete glucose, amino acids, salts, and other nutrients that microorganisms can use to metabolize and grow (28). Sebaceous glands produce triglycerides, free fatty acids, and waxes that also provide additional nutrients for metabolism and growth (28). The ability to metabolize both sweat and sebum nutrients is therefore hypothesized to be key for growth and long-term persistence on skin. An understudied aspect of *C. auris* research is the metabolic strategies it employs to grow in various sweat and sebum nutrient sources.

Previous work has shown that to metabolize glucose *C. albicans* uses conventional carbon metabolism pathways such as glycolysis, the citric acid cycle, and the electron transport chain as well as oxidative phosphorylation for growth (29). However, in nutrient poor environments without glucose *C. albicans* must scavenge and use alternative carbon metabolism (30). For example, *C. albicans* employs β-oxidation (*POX1, FOX2, POT1*), the glyoxylate cycle (*ICL1, MLS1*), and gluconeogenesis to generate glucose from fatty acids for growth (31-36). *C. albicans* also employs the carnitine shuttle (*YAT1, YAT2, CAT2*) to assist in transport of acetyl-CoA (29), multiple secreted lipases (*LIP*) for nutrient acquisition of lipids and adaptation (37, 38), phospholipases and proteases for nutrient acquisition, and amino acid permeases and transporters for additional nutrient uptake (39). Taken together, both conventional and alternative carbon metabolism allow *C. albicans* to grow in various nutrient sources.

Here, we investigate whether *C. auris* employs similar metabolic strategies as *C. albicans* to grow in various nutrient sources like sweat and sebum nutrients. Our work revealed that *C. auris* uses conventional carbon metabolism enzymes and metabolic pathways to metabolize sweat nutrients like glucose, amino acids, and salts. We show that as those nutrients are depleted, *C. auris* changes its metabolic profile and uses alternative carbon metabolism pathways to metabolizes sebum nutrients like fatty acids. Collectively, our results show that *C. auris* shifts its metabolic profile to allow it to grow and persist with limited nutrients in a skin-like environment.

## RESULTS

### *C. auris* growth in rich media and skin-like media

Past studies have shown that *C. auris* is capable of growing in formulations of synthetic sweat medium (20-22). However, no studies have evaluated what genes and metabolic pathways *C. auris* might use to metabolize synthetic sweat medium nutrients, including the addition of a sebum.

To establish this experimental system, we first sought to assess differential growth patterns of *C. auris* AR0387 (Clade I), AR0381 (Clade II), AR0383 (Clade III), and AR0385 (Clade IV), and *C. albicans* SC5314 in nutrient-rich media (YPD) as well as artificial skin-like media supplemented with sebum (Sweat + 0.1% Sebum) (Fig. 1A). In YPD, the area under the curve (AUC) for *C. albicans* compared to each of the *C. auris* clades was significantly higher indicating more robust growth in nutrient-rich conditions (*P* < 0.05) (Fig. 1B). In contrast, all *C. auris* clades had significantly greater AUC than *C. albicans* in Sweat + 0.1% Sebum (*P* < 0.001) (Fig. 1B). Thus, *C. auris* grew better than *C. albicans* in the media simulating skin-like environmental conditions.

**FIG 1.**
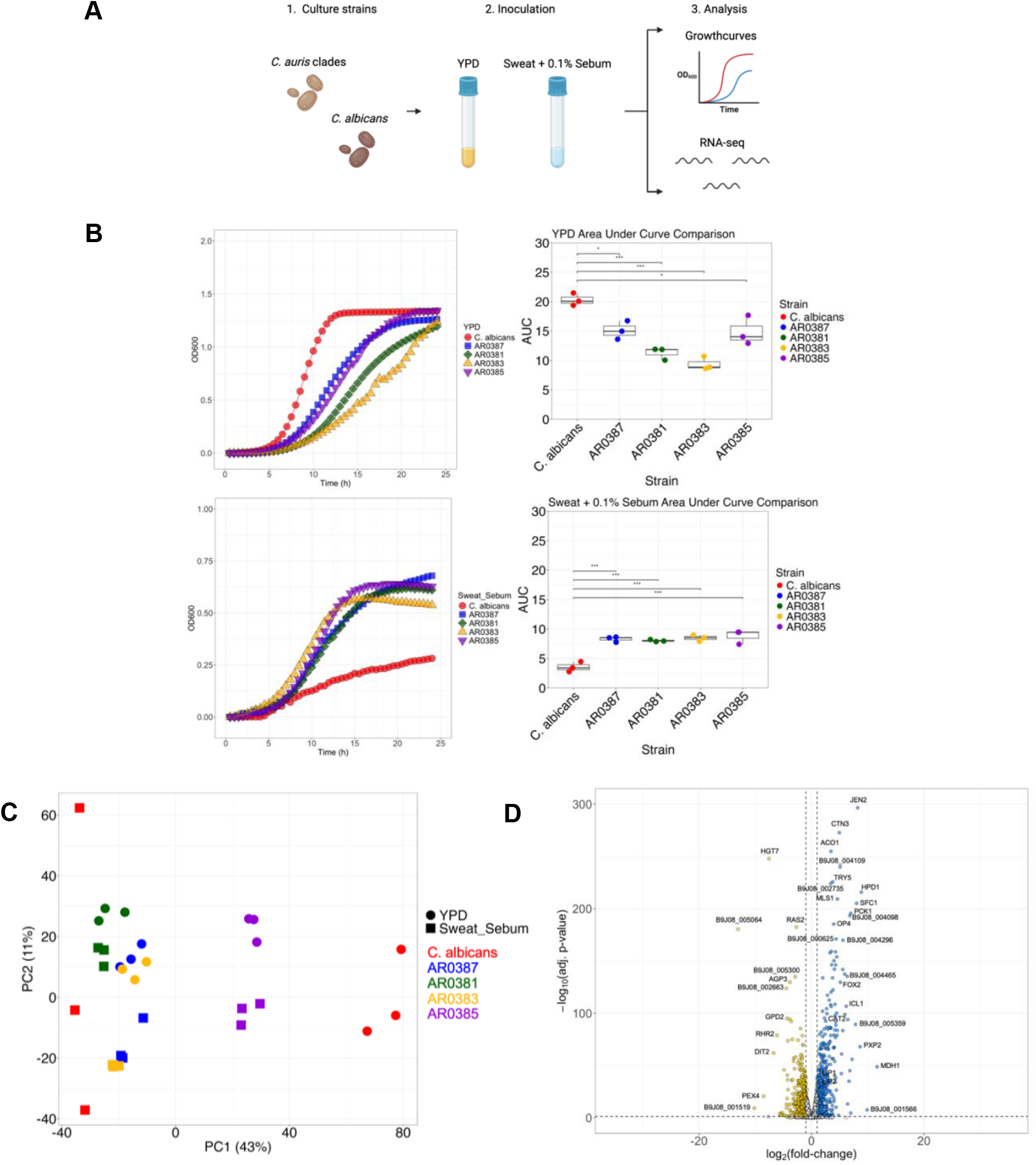
*Candida auris* growth in rich media and skin-like media. (A) *C. albicans* SC5314 and *C. auris* AR0387 (Clade I), AR0381 (Cade II), AR0383 (Clade III), and AR0385 (Clade IV) are cultured in two nutrient sources: Yeast Extract Peptone Dextrose (YPD) and Sweat media supplemented with 0.1% Sebum (Sweat + 0.1% Sebum). Each strain is grown at 34°C, 200 RPM, overnight. Growth analysis is performed on each of the strains and RNA is extracted and sequenced. (B) Each strain is grown in a plate reader at 34°C, 200 RPM, for 24 hr. in YPD and Sweat + 0.1% Sebum media and average data is plotted in a growthcurve. AUC (area under the curve) was calculated and compared between each of the strains. (C) Principal component analysis plot of gene expression for each strain across the different media types using the Z score normalized data. (D) Volcano plot of gene expression between YPD vs. Sweat + 0.1% Sebum for AR0387. Each circle represents a gene expressed per plot. White: no significant difference, Gold: increased in YPD, Blue: increased in Sweat + 0.1% Sebum. Statistics were calculated using one-way ANOVA with Tukey’s post-hoc tests, as appropriate. n = 3, three independent experiments. *p < 0.05, **p < 0.01, ***p < 0.001. Images created with BioRender.com.

Then we sought to understand what genes *C. auris* and *C. albicans* might be using to grow at logarithmic growth phase (8 hr.) in Sweat + 0.1% Sebum compared to YPD using RNA-sequencing. The principal component (PC) analysis (PCA) plot showed that *C. albicans* shifted along PC1 between the different media, while all four clades of *C. auris* shifted along PC2 between the media (Fig. 1C). We also compared growth in the two media types and discovered 620 genes were significantly upregulated by *C. auris* in Sweat + 0.1% Sebum compared to 490 genes significantly upregulated in YPD (Fig. 1D). KEGG pathway analysis also revealed that the genes upregulated in Sweat + 0.1% Sebum were most frequently involved in pathways related to metabolism, biosynthesis of secondary metabolites, peroxisomes, carbon metabolism, and oxidative phosphorylation.

### Growth analysis of *C. auris* strains with deletion of individual lipid metabolism genes

To test our functional hypothesis, we targeted the deletion of differentially expressed genes to assess whether they would provide selective growth defect in Sweat + 0.1% rather than YPD. In our RNA-sequencing analysis we noticed that *C. auris* was upregulating 32 lipid metabolism genes in Sweat + 0.1% Sebum compared to YPD. To interrogate if these genes were key for *C. auris* growth in Sweat + 0.1% Sebum, we made individual mutant strains with Fusion PCR by replacing the genes with a drug resistance marker (40). The genes of interest were *LIP1* (B9J08_004173), *LIP2* (B9J08_004176), *FOX2* (B9J08_002847), *CAT2* (B9J08_000010), and *ICL1* (B9J08_003374). We were specifically interested in the *LIP* genes, because of their ability to metabolize triglycerides (38). We were interested in *FOX2* because of its role in alternative carbon metabolism to metabolize fatty acids through β-oxidation (41). We were interested in *CAT2* because of its ability to assist in transport and break down of acetyl-Coa (29). Lastly, we were interested in *ICL1* because of its role in alternative carbon metabolism to generate citric acid cycle intermediates through the glyoxylate cycle (32-36).

The *lip1Δ, lip2Δ, fox2Δ, cat2Δ*, and *icl1Δ* mutants did not show a growth defect in the nutrient-rich YPD compared to *C. auris* WT and the AUC was not significantly different between any of the strains (*P* > 0.05) (Fig. 2A). For the Sweat + 0.1% Sebum curves, the *LIP* mutants grew equally as well as *C. auris* WT (Fig. 2A). However, *fox2Δ, cat2Δ*, and *icl1Δ* had notable growth defects in Sweat + 0.1% Sebum and the AUC was significantly lower than *C. auris* WT (*P* < 0.01) (Fig. 2A). Collectively these results indicate that *fox2Δ, cat2Δ*, and *icl1Δ* had significantly abrogated growth in Sweat + 0.1% Sebum media, but not in YPD media.

**FIG 2.**
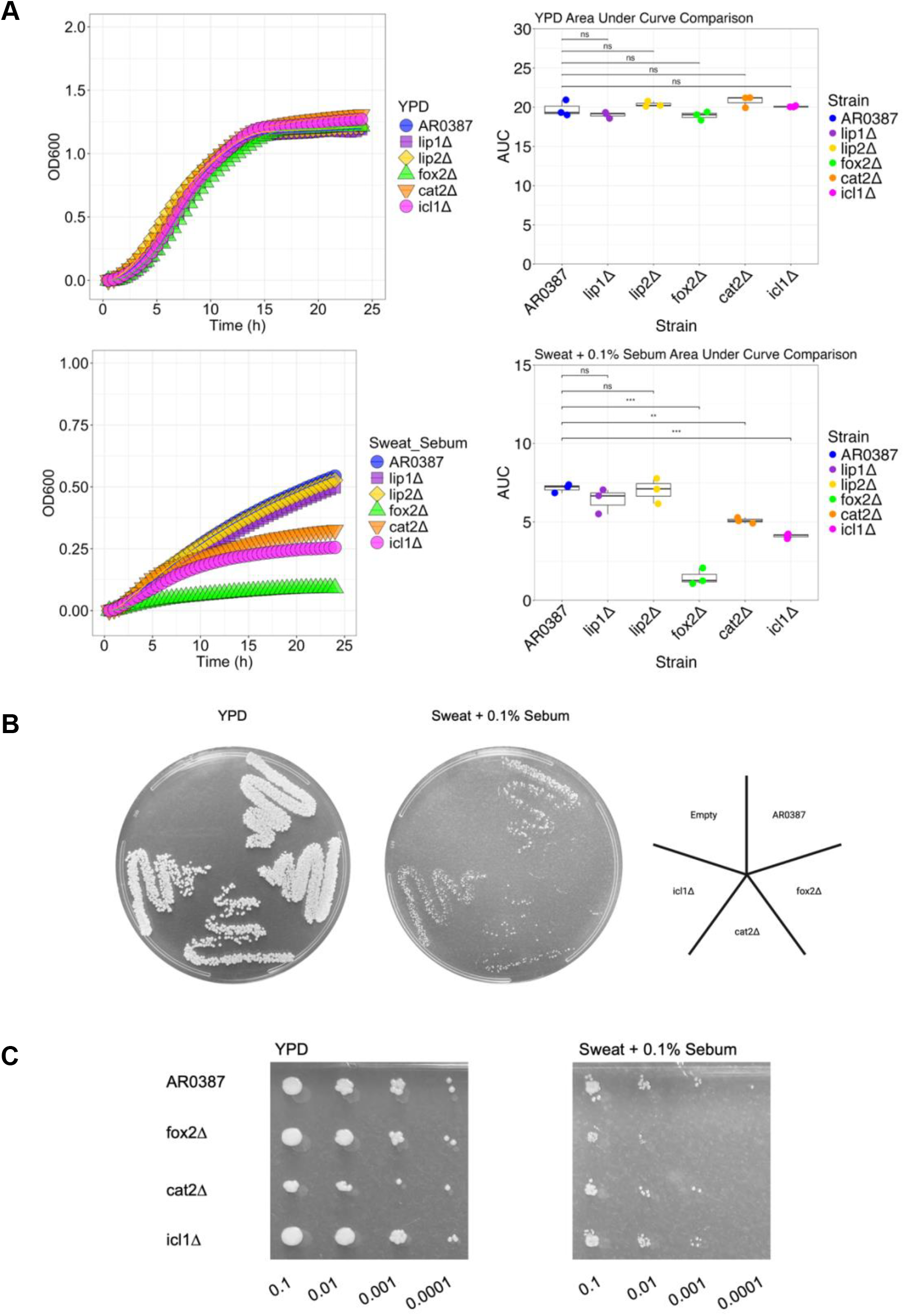
Growth analysis of *C. auris* strains with deletion of individual lipid metabolism genes. (A) Each strain is grown in a plate reader at 34°C, 200 RPM, for 24 hr. in YPD and Sweat + 0.1% Sebum media and average data is plotted in a growthcurve. AUC (area under the curve) was calculated and compared between each of the strains. (B) Strains were cultured in YPD and Sweat + 0.1% Sebum, set to an OD of 0.1, and swabbed on their respective agar. (C) Strains were cultured in YPD and Sweat + 0.1% Sebum, set to an OD of 0.1, and serial diluted on their respective agar. Statistics were calculated using one-way ANOVA with Tukey’s post-hoc tests, as appropriate. n = 3, three independent experiments. *p < 0.05, **p < 0.01, ***p < 0.001. Images created with BioRender.com.

Next, to complement the *in vitro* broth media assessment, we assessed if growth defects were observed on solid agar. Again, no growth defects were observed on YPD agar (Fig. 2B). However, on the Sweat + 0.1% Sebum agar growth was observed to be abrogated for *fox2Δ, cat2Δ*, and *icl1Δ* (Fig. 2B). To assess the growth capacity of *C. auris* WT and *fox2Δ, cat2Δ*, and *icl1Δ* we then performed a serial dilution spot assay onto YPD or Sweat + 0.1% Sebum agar. Again, growth defects were observed specifically on Sweat + 0.1% Sebum (Fig. 2C). Collectively, *fox2Δ, cat2Δ*, and *icl1Δ* displayed abrogated growth defects on Sweat + 0.1% Sebum agar, but not YPD agar.

### Sweat and sebum gradient assay plates demonstrate nutrient preferences between *C. auris* WT and mutant strains

We next assayed whether different growth phenotypes were observable for *C. auris* WT and the mutant strains when sweat and sebum nutrients were varied. To accomplish this, a two-dimensional gradient assay plate was used to grow wild type and mutant strains at eight concentrations of sweat (0.03X to 4X) and sebum (0.002% to 0.25%) for a total of sixty-four growth conditions. Each gradient assay plate was inoculated and assessed for growth at 24 and 48 hours. Data were normalized and optical density was plotted as a heatmap to show differences in nutrient preferences (Fig. 3A).

**FIG 3.**
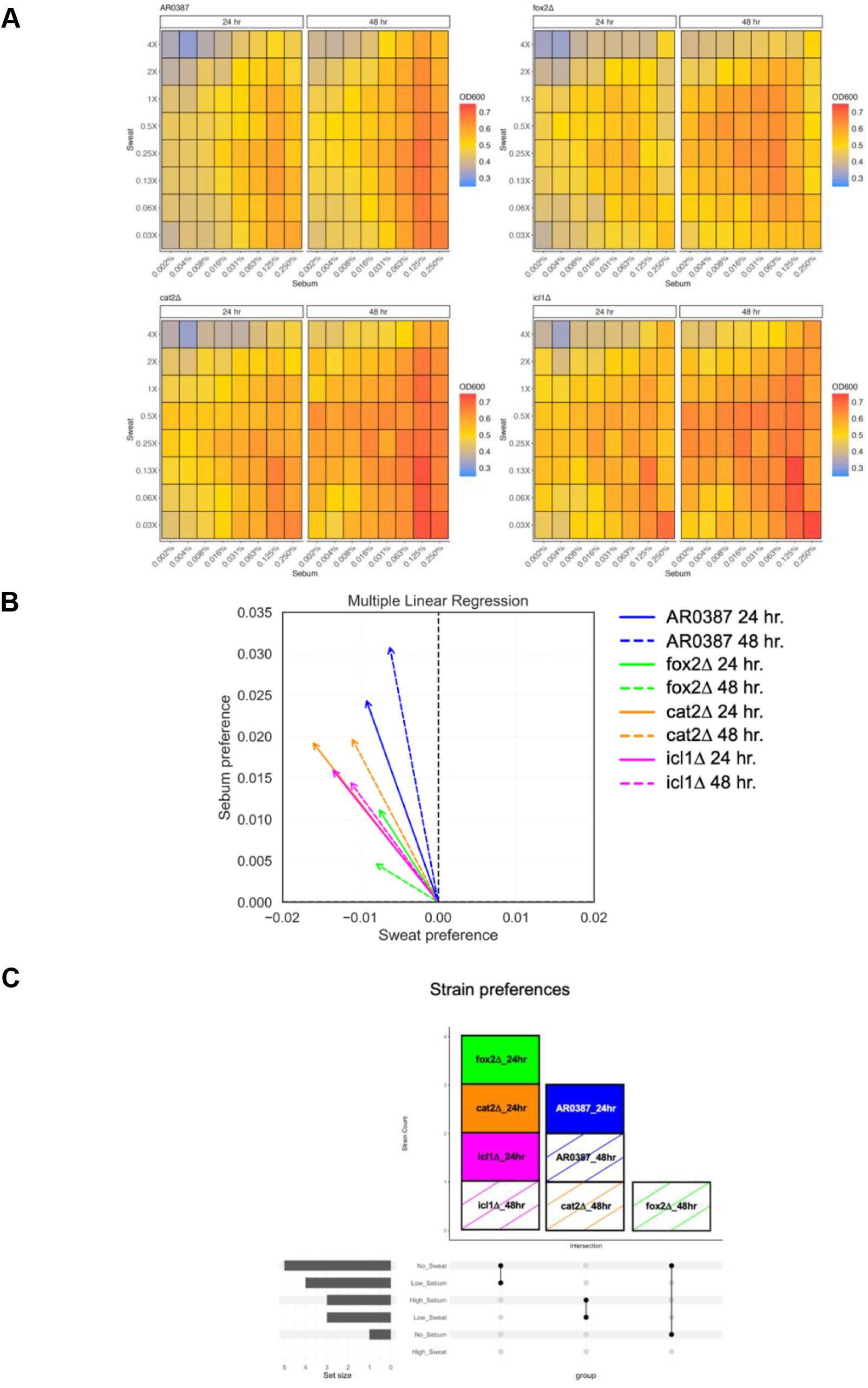
Sweat and sebum gradient assay plates demonstrate nutrient preferences between *C. auris* WT and mutant strains. (A) AR0387 and mutant strains are cultured at 34°C, 200 RPM overnight in Sweat + 0.1% Sebum. Cells are washed, set to an OD600 of 0.1, and inoculated in a gradient assay plate of Sweat (0.03X, 0.06X, 0.13X, 0.25X, 0.5X, 1X, 2X, 4X) and Sebum (0.002%, 0.004%, 0.008%, 0.016%, 0.031%, 0.063%, 0.125%, 0.25%) at 34°C, 200 RPM and grown for 24 and 48 hours. OD600 was measured at each time point. (B) Multiple linear regression analysis of growth in gradient assay plates. Analysis generated using Z score normalized data. (C) Upset plot of strain preferences based on 6 nutrient conditions in each environment: High Sweat, Low Sweat, No Sweat, High Sebum, Low Sebum, No Sebum.

Next, a multiple linear regression analysis was performed for *C. auris* WT and the mutant strains to determine nutrient preferences for growth in the gradient assay plate over time. *C. auris* WT had a sebum preference over time. *Fox2Δ* had a slight sebum preference at 24 hours, but lost preferences over time likely due to the inability to metabolize sebum nutrients without *FOX2. Cat2Δ* also had a slight sebum preference at 24 hours, but developed a stronger sebum preference over time likely because it can still metabolize sebum nutrients without *CAT2. Icl1Δ* had a slight sebum preference over time, but most likely does not gain a strong sebum preference due to the inability to adequately metabolize sebum nutrients without *ICL1*. Collectively, these results indicate that *fox2Δ* and *icl1Δ* had the most distinct nutrient preferences, whereas *cat2Δ* behaved more similarly to *C. auris* WT (Fig. 3B; Fig. 3C).

### RNA-sequencing analysis of *C. auris* WT compared to mutant strains

Next, we wanted to explore the genetic pathways altered within *fox2Δ, cat2Δ*, and *icl1Δ* that accompany the previously observed growth defects. To accomplish this, we assessed differences in gene expression at logarithmic growth phase (8 hr.) in Sweat + 0.1% Sebum by comparing *C. auris* WT and *fox2Δ, cat2Δ*, and *icl1Δ* with RNA-sequencing analysis.

Volcano plots were generated to identify the most differentially expressed genes between *C. auris* WT and *fox2Δ, cat2Δ*, and *icl1Δ* with a log2 fold change > 1 and adjusted P-value < 0.05. (Fig. 4A). As a control, and reassuringly, the most differentially expressed gene for each comparison was the gene removed from the genome in each mutant strain. When grown in Sweat + 0.1% Sebum, *fox2Δ* upregulated 184 genes, *cat2Δ* upregulated 40 genes, and *icl1Δ* upregulated 67 genes compared to the wild type. Sixteen genes were commonly upregulated and 3 genes were commonly downregulated by all three mutants in comparison to *C. auris* WT (Fig. 4B). Of the 16 commonly upregulated genes, 7 were annotated: *ILV3, LEU1, MET16, PRX1, ERG24, ARG3*, and *DAG7*. Of the 3 commonly downregulated genes 2 were annotated: *WOR4* and *AQY1*. KEGG pathway analysis revealed that the upregulated genes were primarily involved in biosynthesis of secondary metabolites and amino acid synthesis whereas the downregulated genes did not have sufficient information for analysis.

**FIG 4.**
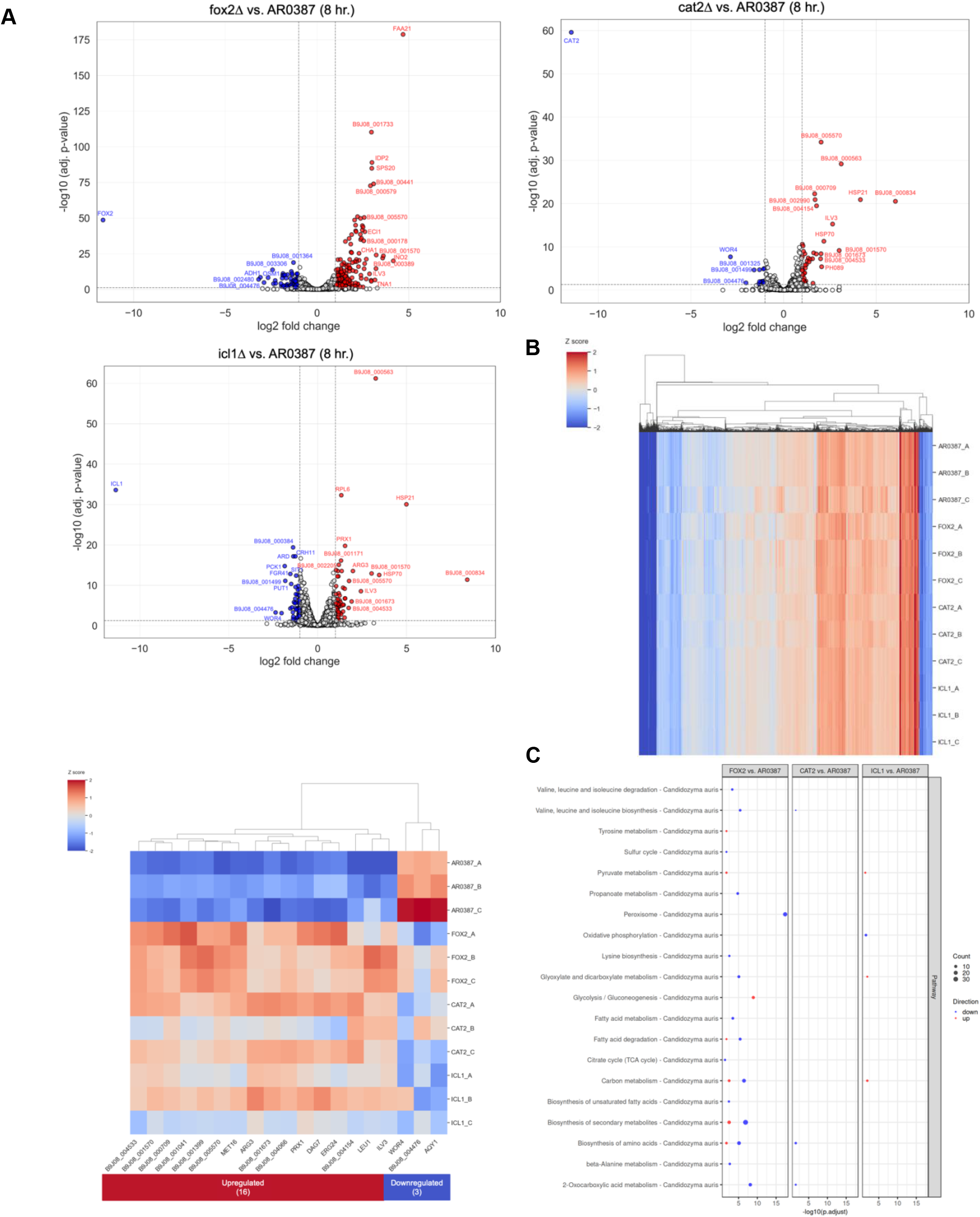
RNA-sequencing analysis of *C. auris* WT compared to mutant strains (A) Volcano plots of gene expression between AR0387. Each circle represents a gene. White: no significant difference, Blue, downregulated, Red: upregulated. (B) Heatmap of AR0387 and mutant strains gene expression in Sweat + 0.1% Sebum using the Z score normalized data. Additional heatmap of genes commonly upregulated and downregulated by *fox2Δ, cat2Δ*, and *icl1Δ* compared to AR0387. (C) Pathway analysis of AR0387 and mutant strains gene expression in Sweat + 0.1% Sebum.

Pathway analysis was also done to evaluate which pathways were upregulated and downregulated comparing each mutant to *C. auris* WT (Fig. 4C). For *fox2Δ*, we observed increased expression of tyrosine metabolism, pyruvate metabolism, and glycolysis and gluconeogenesis genes (Fig. 4C). For *fox2Δ*, we also observed decreased expression of valine, leucine, and isoleucine degradation and biosynthesis, sulfur cycle, propanoate metabolism, peroxisome, lysine biosynthesis, glyoxylate and dicarboxylate metabolism, fatty acid metabolism and degradation, citrate cycle (TCA cycle), carbon metabolism, biosynthesis of unsaturated fatty acids and secondary metabolites, biosynthesis of amino acids, beta-Alanine metabolism, and 2-oxocarboxylic acid metabolism. These results suggest that when *FOX2* is deleted the cell responds by increasing the expression of multiple pathways to metabolize available sweat nutrients to account for its hindered ability to metabolize sebum nutrients.

For *cat2Δ*, we observed decreased expression of valine, leucine, and isoleucine biosynthesis, biosynthesis of amino acids, and 2-oxocarboxylic acid metabolism (Fig. 4C). These results suggest that when *CAT2* is deleted it does not affect gene expression as significantly and metabolism is not as altered.

For *icl1Δ*, we observed increased expression of pyruvate metabolism, glyoxylate and dicarboxylate metabolism, and carbon metabolism. *Icl1Δ* also had decreased expression of oxidative phosphorylation (Fig. 4C). These results suggest that like *FOX2*, when *ICL1* is deleted the cell responded by increasing the expression of multiple pathways to metabolize available sweat nutrients to account for its hindered ability to metabolize sebum nutrients.

Collectively, these results suggest that *FOX2, CAT2*, and *ICL1* are important for growth in Sweat and Sebum. Moreover, upregulation of additional genes and pathways sheds light on how metabolism occurs in a variable sweat and sebum environment.

## DISCUSSION

*C. auris* is a fungal pathogen of increasingly urgent health risk due to multidrug-resistance and its ability to cause hard to treat bloodstream infections. However, skin is the primary site of *C. auris* colonization and this predisposes patients to shedding of the organism which promotes seeding of the environment and transmission. Understanding how *C. auris* can grow on skin and what nutrients it might be utilizing to do so is a significant gap in knowledge regarding this pathogen. Our data reveal that encoding diverse conventional and alternative carbon metabolism genes facilitates growth and nutrient utilization in a skin-like environment.

Skin is a nutritionally diverse location and has four distinct microenvironments (sebaceous, dry, moist, and foot) with varied density of sweat and sebaceous glands depending on body site (27). Sweat glands provide glucose, amino acids, salts, and other nutrients and sebaceous glands provide triglycerides, fatty acids, and waxes (28). However, these nutrients are not abundant on skin, so microorganisms that grow there are generally adapted to this low nutrient environment.

The healthy skin mycobiome, or the collection of fungi on skin, is dominated by *Malassezia* species (42, 43). *Malassezia* can grow across the entire body surface, however they are lipophilic, or “fat-loving”, and prefer the lipid-rich environment of the hair follicles/sebaceous gland where there is sebum (43). This is because *Malassezia* lack fatty acid synthase and cannot synthesize fatty acids on their own, so they scavenge them from their environment (44) Consequently, *Malassezia* have lipases, phospholipases, and sphingomyelinases (45-47) to break down extracellular lipids from sebum to fatty acids, allowing them to grow (30, 43). Malassezia also lack carbon metabolism genes which is likely an evolutionary consequence of their long-term colonization of the sebum rich areas of skin (44).

*C. albicans*, which is an ascomycete genetically more similar to *C. auris* than *Malassezia* species, can grow on skin but has a greater tropism for the gastrointestinal flora, oral cavity, and reproductive tract. *C. albicans* also differs from *Malassezia* because it employs its own set of strategies for metabolism in various nutritional environments. For example, if glucose, the preferred carbon source, is present it will be metabolized with conventional carbon metabolism pathways such as glycolysis to form acetyl-CoA. Then acetyl-CoA will enter the citric acid cycle, a conventional carbon metabolism pathway, to produce eight intermediates and multiple coenzymes. Two coenzymes, NADH and FADH2, enter the electron transport chain and generate ATP through oxidative phosphorylation allowing cellular growth (33). If glucose is unavailable or in low quantities, *C. albicans* uses alternative carbon metabolism pathways to grow instead. One such pathway is β-oxidation, which is carried out by three enzymes (*POX1, FOX2, POT1*) in four metabolic steps (29). In β-oxidation, fatty acids are broken down into acetyl-CoA (29). Then acetyl-CoA can enter the glyoxylate cycle, another alternative carbon metabolism pathway, which is unique to *Candida* species, bacteria, and plants (33). In this process, isocitrate is hydrolyzed to glyoxylate by *ICL1*. Then acetyl-CoA, from β-oxidation, is condensed with the previously produced glyoxylate to produce malate by *MLS1*. Malate, a citric acid cycle intermediate, is converted to oxaloacetate, then citrate, and back to isocitrate. This process replenishes intermediates of the citric acid cycle allowing it to function when nutrients are low.

Additionally, the oxaloacetate from the glyoxylate cycle can enter gluconeogenesis, another alternative carbon metabolism pathway, to produce glucose which provides energy to sustain the cell. *C. albicans* also uses the carnitine shuttle (*YAT1, YAT2, CAT2*) to transport acetyl-CoA into the mitochondria and peroxisome for downstream metabolism (29). *C. albicans* also has many secreted lipases (*LIP*) at its disposal for additional metabolism of lipids too (37, 38). Overall, *C. albicans* has many strategies it can use to metabolize nutrients in either rich or poor environments to grow with conventional or alternative carbon metabolism respectively.

Other *Candida species* also use alternative carbon metabolism when glucose is not available. For example, *Candida lusitaniae* uses β-oxidation (*FOX2*) and the glyoxylate cycle (*ICL1*) to metabolize fatty acids as a carbon source to grow and transports acetyl-units with carnitine acetyl-transferase systems (*CAT2*) (48). *Candida tropicalis* uses β-oxidation (*FOX2*) to break down short length fatty acids to generate acetyl-CoA and then transports it to the mitochondria and peroxisome via carnitine acetyl-transferase systems (*CAT2*) (49). *Candida glabrata* uses β-oxidation (*FOX2*) to generate acetyl-CoA, carnitine acetyl-transferase systems (*CAT2*) to move acetyl-CoA, and the glyoxylate cycle (*ICL1*) to process acetyl-CoA (50).

Prior to this study, there has been limited knowledge about what metabolic pathways *C. auris* might use to grow in a skin-like environment. Here, we showed that *C. auris* uses lipid metabolism genes to grow in Sweat + 0.1% Sebum media, but those genes are not essential for growth in glucose rich YPD. We found that key genes that facilitate this growth are *FOX2, CAT2*, and *ICL1*, similar to other *Candida species*.

*FOX2* is important for *C. auris* because it drives the alternative carbon metabolism pathway β-oxidation, which metabolizes fatty acids to acetyl-CoA (Fig. 5A; Fig. 5B; Fig. 5C). Major nutrients of our Sweat + 0.1% Sebum media are fatty acids, not glucose. Therefore, if *FOX2* cannot function, then fatty acids will not be properly metabolized, and the cell will not grow. Similarly, if *CAT2* cannot function then acetyl-CoA will not be properly transported to the mitochondria or peroxisome, downstream metabolic pathways will not function, and the cell will not grow (Fig. 5A; Fig. 5C). Lastly, if *ICL1* cannot function then the glyoxylate cycle will not produce citric acid cycle intermediates and coenzymes properly and the cell will not grow (Fig. 5A. Fig. 5C). This corelates with past studies where a *C. albicans FOX2* mutant (51), an *ICL1* mutant (32), and a *CAT2* mutant (52) did not grow on oleic acid as a nutrient source.

**FIG 5.**
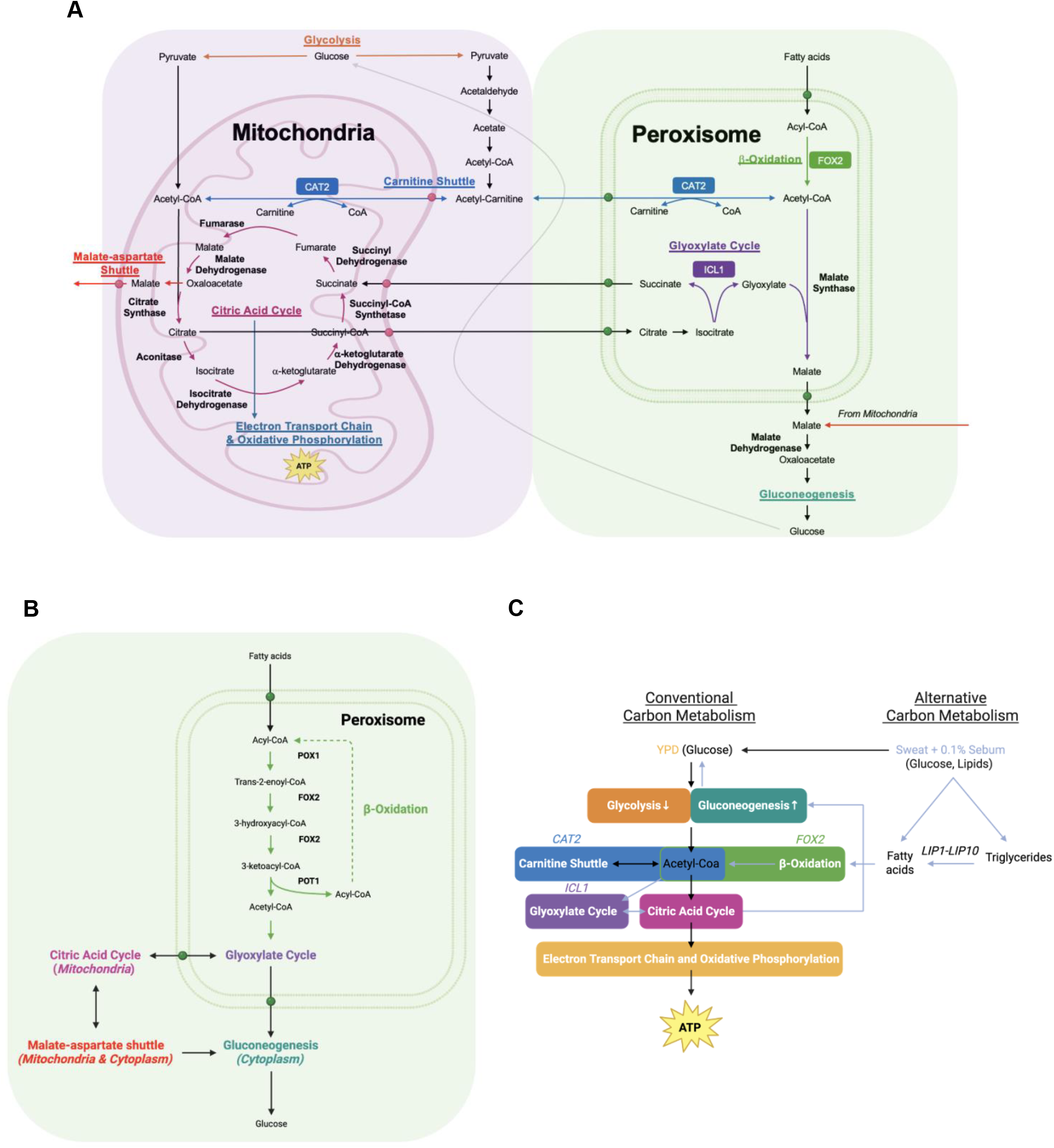
Metabolism in *C. auris* in the mitochondria, peroxisome, and cytosol. (A) Model for metabolism in the mitochondria, peroxisome, and cytosol. Pathways shown are glycolysis, citric acid cycle, electron transport chain and oxidative phosphorylation, carnitine shuttle, β-oxidation, glyoxylate cycle, malate-aspartate shuttle, and gluconeogenesis. (B) β-oxidation. (C) Simplified metabolism model. Images created with BioRender.com.

We were also interested to learn if *C. auris* had growth preferences in Sweat + 0.1% Sebum. When we grew *C. auris* in a gradient assay plate with a broad range of nutrients we noticed that it could grow in the sweat nutrients (glucose), but it preferred the sebum nutrients (fatty acids) (Fig. 3). This was noteworthy because it indicated that *C. auris* metabolized the sweat nutrients with conventional carbon metabolism first, and then the sebum nutrients with alternative carbon metabolism second. This long-term sebum preference is similar to skin commensal *Malassezia* species which are lipophilic, or ‘fat-loving’, and prefer the lipid-rich environment of the hair follicles/sebaceous gland (43). This could explain why *C. auris* eventually navigates into the pilosebaceous unit where there is available sebum (22, 23). Perhaps, *C. auris* is also a ‘fat-loving’ microbe too which could explain why it can grow and persist on skin so effectively. Moreover, *C. auris* may persist in this niche unlike other pathogens because the available fatty acids aren’t antimicrobial, but rather a source of nutrition (44). *C. auris* may perhaps outcompete *Malassezia* in this niche. *Fox2Δ* did not metabolize the range of available sebum nutrients like the wild type validating its importance in fatty acid metabolism. Curiously, *cat2Δ* behaved more similarly to the wild type. Perhaps this is because the carnitine shuttle is not essential for alternative carbon metabolism, and *C. auris* might have other ways to move acetyl units into the organelles like *C. albicans* (29). *Icl1Δ* never developed a strong sebum preference relative to the wild type. This also indicated *ICL1*’s importance in fatty acid metabolism.

We also discovered that *fox2Δ* upregulated genes in glycolysis/gluconeogenesis and pyruvate metabolism pathways compared to the wild type in Sweat + 0.1% Sebum (Fig. 4C). We hypothesize this is occurring because *fox2Δ* is trying to metabolize the glucose and other sweat nutrients, but not the fatty acids in sebum, since it is unable to do so without the gene. *Fox2Δ* downregulated sixteen metabolic pathways (Fig. 4C).

We hypothesize this is occurring because *FOX2* is a key alternative carbon metabolism gene and without it downstream metabolism is significantly altered in many pathways. *Cat2Δ* only downregulated three metabolic pathways (Fig. 4C). We hypothesize this is occurring because *CAT2* is not as directly involved in alternative carbon metabolism. Like *fox2Δ, icl1Δ* upregulated pyruvate metabolism as well as glyoxylate, dicarboxylate, and carbon metabolism (Fig. 4C). We hypothesize this is occurring because *icl1Δ* is also trying to metabolize glucose from sweat nutrients because it cannot metabolize the fatty acids in sebum (Fig. 4C). Collectively, these results suggest that *FOX2* and *ICL1* are important for growth in Sweat + 0.1% Sebum media with *CAT2* playing a minor role.

We also discovered 16 commonly upregulated and 3 commonly downregulated genes for the mutants compared to the wild type (Fig. 4B). Of the 16 commonly upregulated genes, 7 had annotations. *ILV3, LEU1, MET16*, and *ARG3* were involved in amino acid biosynthesis. *PRX1* is involved with reducing hydrogen peroxide to water, *DAG7* is a possible secretory protein, and *ERG24* is involved with ergosterol biosynthesis. Perhaps the mutants are upregulating multiple amino acid biosynthesis genes because the amino acids they synthesize (valine, isoleucine, leucine, sulfur amino acids, and arginine) are important for proper metabolism in a low nutrient environment like Sweat + 0.1% Sebum. *PRX1* may be upregulated to mitigate hydrogen peroxide production during metabolism. Lastly, *ERG24* may be upregulated for proper synthesis of ergosterol in Sweat + 0.1% Sebum. Of this group of genes, it is possible that *ERG24* could be a potential drug target as *C. albicans* mutants of this gene are susceptible to allylamine antifungals (terbinafine) as well as cellular inhibitors including cycloheximide, cerulenin, fluphenazine, and brefeldin A (53). Of the 3 commonly downregulated genes 2 were annotated. These included *WOR4*: has domain(s) with predicted role in regulation of DNA-templated transcription and *AQY1*: Ortholog(s) have water channel activity. It is unclear why these genes were commonly downregulated as they were not related to metabolism. A limitation of this study was that the additional genes discovered could not be analyzed because of insufficient annotations in the KEGG pathway database. More work will be done in the future to understand why the additional differentially expressed genes may be relevant to sweat and sebum metabolism.

Overall, we showed that *C. auris* has more robust growth than *C. albicans* in Sweat + 0.1% Sebum media. We also showed that conventional carbon metabolism facilitates growth in sweat nutrients when glucose is present. Alternative carbon metabolism facilitates growth in sebum when fatty acids are present requiring genes like *FOX2, CAT2*, and *ICL1*. Additionally, we found more genes and metabolic pathways to explore to better understand *C. auris* growth capabilities. We hope these efforts will pave the way for better understanding of *C. auris* metabolism and advance understanding of the genes and pathways that could serve as potential targets for control. More effective strategies are needed to mitigate consequences of *C. auris* on skin to mitigate risks for transmission and development of bloodstream infection.

## MATERIALS AND METHODS

### STRAINS

The *C. auris* and *C. albicans* strains used in this study are listed in Table S1 in the supplemental material.

### MEDIA AND GROWTH CONDITIONS

Strains were grown and plated in two different nutrient sources. The first nutrient source was YPD (Yeast extract peptone dextrose) Broth (Sigma-Aldrich) and YPD Broth + Agar (Sigma-Aldrich) (2%). The second nutrient source was Sweat + 0.1% Sebum with commercial Sweat + 0.1% Sebum used to collect initial data in (Fig. 1). This formulation consisted of three components: Artificial Eccrine Perspiration (Pickering Laboratories, Cat number: 1700-0023), Artificial Sebum (Pickering Laboratories, Cat number: 1700-0700), and 1% Tween 80 (MP Biomedicals, Cat number: 103170). The second Sweat + 0.1% Sebum formulation was a defined media used to collect data in (Fig. 2; Fig. 3; Fig. 4) adopted from Swaney et al., 2023 (27). This formulation consisted of three components: Basal media, Artificial Sweat, and Artificial Sebum. The contents of the media can be found in Table S2. Sweat + 0.1% Sebum agar plates were made by adding 1.5% agar to the Sweat + 0.1% Sebum media previously described.

### GROWTH CURVES AND GROWTH METRIC ANALYSIS

Pure culture isolates of the *C. albicans* and *C. auris* wild type and mutant strains were grown initially at 34ºC, 200 RPM, overnight in YPD or Sweat + 0.1% Sebum. After normalizing the initial inoculum to OD600 0.1, 10 μl of the freshly washed cells were added to 190 μl of media in a 96-well plate (Nunc™ Edge™ 96-Well, Non-Treated, Flat-Bottom Microplate (Thermo Fisher, Cat: 267544). The plates were placed in a plate reader (Agilent BioTek Epoch 2 Microplate Spectrophotometer, Fisher Scientific) at 34ºC, 200 RPM, for 24 hours. The Growthcurver script was adapted from Sprouffske and Wagner 2016 (54) to plot growth curves and calculate growth metrics such as AUC (area under the curve), k (carrying capacity), r (growth rate), and t_mid (inflection point of the curve). Statistics were calculated using one-way ANOVA with Tukey’s post-hoc tests in R.

### GENE DELETION STRATEGY

Mutant strains were generated from *C. auris* AR0387 with the Fusion PCR protocol and were confirmed with the Colony PCR protocol adopted from Schwarzmuller et al., 2014 (40). Sequencing primers were also generated upstream of the gene of interest that was deleted, in the middle of the selectable marker, and at the 3’ end of the selectable marker to ensure proper deletion of the gene. All primers are available in Table S3.

### SWAB AND SERIAL DILUTION PLATES

*C. auris* AR0387 and mutant strains were grown at 34ºC, 200 RPM, overnight in YPD or Sweat + 0.1% Sebum. After normalizing the initial inoculum to OD600 0.1, swabs were inoculated and streaked on corresponding YPD agar or Sweat + 0.1% Sebum agar, incubated at 34ºC for 24 hours, and imaged.

Then the normalized inoculum was serial diluted 10-fold, twelve times into a 96-well plate. Then a replicator (Boekel, 96-Pin Microplate Replicator, 140500) was used to transfer 1 μl of cells to a corresponding YPD or Sweat + 0.1% Sebum agar plate, incubated at 34ºC for 24 hours, and imaged.

### GRADIENT ASSAY PLATES

A gradient assay plate of sweat and sebum was generated which was adopted from Swaney, et al., 2023 (27). In this plate, there are eight concentrations of sweat (0.03X, 0.06X, 0.13X, 0.25X, 0.5X, 1X, 2X, and 4X) and eight concentrations of sebum (0.002%, 0.004%, 0.008%, 0.016%, 0.031%, 0.063%, 0.125%, 0.25%) for a total of sixty-four growth conditions. The remaining wells of the plate were used as controls. Column 9 was the last sweat dilution, column 10 was the last sebum dilution, column 11 was basal media, and column 12 was YPD. Well H12 was also inoculated with *C. auris* AR0387 as a positive control. Gradient assay plates were grown at 34ºC, 200 RPM. OD600 values were collected with a plate reader at 0 hours, 24 hours, and 48 hours.

Then, the log2 of the sweat and sebum concentrations were used as feature X and the OD600 was used at feature Y. Then a linear regression was trained for each strain. A slope was generated from the X and Y coefficients at each time point to determine how growth changed in sweat and sebum.

Slope thresholds were used to generate nutrient preferences. High sweat = 0° to 60°, Low sweat = 60° to 120°, and No sweat = 120° to 180°. High sebum = 60°-120°, Low sebum = 30° to 60° and 120° to 150°, and No Sebum = 0° to 30° and 150° to 180°.

### RNA SEQUENCING AND ANALYSIS

*C. albicans* and *C. auris* wild type strains and mutant strains were grown at 34ºC, 200 RPM overnight for 8 and 15 hours in 5 ml of YPD or Sweat + 0.1% Sebum in triplicate. RNA-sequencing reads were mapped using STAR v2.7.11b (55) to the *C. albicans* SC5314 genome (GCF_000182965.3) from NCBI RefSeq. HOMER v5.1 (56) was used to create tag directories based on mapped data with the script “makeTagDirectory - format sam -checkGC -sspe”, and to calculate raw counts and TPM (transcripts per million) values using “analyzeRepeats.pl rna -count exons”. Differentially expressed genes were identified using HOMER script “getDiffExpression.pl -AvsA -repeats”, which implements DESeq2 (57), with significance thresholds of log2 fold change > 1 and adjusted *P* value < 0.05. Volcano plots and heat maps were generated to visualize expression difference between *C. auris* wild type and mutant strains. KEGG pathway enrichment analysis was performed on differentially expressed genes using R package ClusterProfiler v4.10.1 (58) with an adjusted *P* value threshold of < 0.05.

### STATISTICAL ANALYSIS

All experiments were performed with at least three biological replicates as indicated in the figure legends. Analyses were conducted with R and python. Statistics were calculated using one-way ANOVA with Tukey’s post-hoc tests in R. *P* values < 0.05 were considered statistically significant.

## ACKNOWLEDGMENTS

We thank the members of our laboratory for critical discussion on experimental design, execution, and analysis. We thank Lukian Robert for assistance in screening mutants, Milan Stolpman for assistance in gradient assay plates set ups, and Diana Proctor for bioinformatic analysis. We thank the Karl Kuchler Lab for providing the pTS50 plasmid used in the Gene Deletion Strategy. We thank Teresa O’Meara’s laboratory for thoughtful discussion on our experimental design and suggestions for the manuscript. We thank the NIH Intramural Sequencing Center (NISC) for sequencing isolates. We thank the Microarray and Single Cell Genomics Core at NIH for quantification of RNA. This study utilized the computational resources of the NIH HPC Biowulf Cluster (http://hpc.nih.gov). J.A.S. is an Associate Fellow of the Canadien Institute for Advanced Research (CIFAR) program Fungal Kingdom: Threats & Opportunities.

## FUNDING

This work was supported by the Division of Intramural Research of the National Human Genome Research Institute (NHGRI).

## DATA AVAILABILITY

*NCBI BioProject to house all the data*

. *BioSamples for the WT RNAseq*.

*BioSamples for the mutant RNAseq*.

## ETHICS APPROVAL

## ADDITIONAL FILES

## SUPPLEMENTAL MATERIAL

Supplemental figures:

Table S1. Strains used and designed in this study.

Table S2. Artificial Sweat, Artificial Sebum, and Basal Media recipes used in this study. Table S3. Primers used in this study.

Figure S1. Volcano plots of gene expression between AR0387 and mutant stains (15 hr.)

Figure S2. Heatmap of AR0387 and mutant strains gene expression in Sweat + 0.1% Sebum using the Z score normalized data (15 hr.)

Figure S3. Pathway analysis of AR0387 and mutant strains gene expression in Sweat + 0.1% Sebum (15 hr.)

## REFERENCES

1. Lockhart SR, Etienne KA, Vallabhaneni S, Farooqi J, Chowdhary A, Govender NP, Colombo AL, Calvo B, Cuomo CA, Desjardins CA, Berkow EL, Castanheira M, Magobo RE, Jabeen K, Asghar RJ, Meis JF, Jackson B, Chiller T, Litvintseva AP. 2017. Simultaneous Emergence of Multidrug-Resistant Candida auris on 3 Continents Confirmed by Whole-Genome Sequencing and Epidemiological Analyses. Clin Infect Dis 64:134–140.

2. Satoh K, Makimura K, Hasumi Y, Nishiyama Y, Uchida K, Yamaguchi H. 2009. Candida auris sp. nov., a novel ascomycetous yeast isolated from the external ear canal of an inpatient in a Japanese hospital. Microbiol Immunol 53:41–4.

3. Spruijtenburg B, Badali H, Abastabar M, Mirhendi H, Khodavaisy S, Sharifisooraki J, Taghizadeh Armaki M, de Groot T, Meis JF. 2022. Confirmation of fifth Candida auris clade by whole genome sequencing. Emerg Microbes Infect 11:2405–2411.

4. Khan T, Faysal NI, Hossain MM, Mah EMS, Haider A, Moon SB, Sen D, Ahmed D, Parnell LA, Jubair M, Chow NA, Chowdhury F, Rahman M. 2024. Emergence of the novel sixth Candida auris Clade VI in Bangladesh. Microbiol Spectr 12:e0354023.

5. Suphavilai C, Ko KKK, Lim KM, Tan MG, Boonsimma P, Chu JJK, Goh SS, Rajandran P, Lee LC, Tan KY, Shaik Ismail BB, Aung MK, Yang Y, Sim JXY, Venkatachalam I, Cherng BPZ, Spruijtenburg B, Chan KS, Oon LLE, Tan AL, Tan YE, Wijaya L, Tan BH, Ling ML, Koh TH, Meis JF, Tsui CKM, Nagarajan N. 2024. Detection and characterisation of a sixth Candida auris clade in Singapore: a genomic and phenotypic study. Lancet Microbe 5:100878.

6. Garcia-Bustos V. 2024. Is Candida auris the first multidrug-resistant fungal zoonosis emerging from climate change? mBio 15:e0014624.

7. Adams E, Quinn M, Tsay S, Poirot E, Chaturvedi S, Southwick K, Greenko J, Fernandez R, Kallen A, Vallabhaneni S, Haley V, Hutton B, Blog D, Lutterloh E, Zucker H, Candida auris Investigation W. 2018. Candida auris in Healthcare Facilities, New York, USA, 2013-2017. Emerg Infect Dis 24:1816–1824.

8. Kumar J, Eilertson B, Cadnum JL, Whitlow CS, Jencson AL, Safdar N, Krein SL, Tanner WD, Mayer J, Samore MH, Donskey CJ. 2019. Environmental Contamination with Candida Species in Multiple Hospitals Including a Tertiary Care Hospital with a Candida auris Outbreak. Pathog Immun 4:260–270.

9. Sexton DJ, Bentz ML, Welsh RM, Derado G, Furin W, Rose LJ, Noble-Wang J, Pacilli M, McPherson TD, Black S, Kemble SK, Herzegh O, Ahmad A, Forsberg K, Jackson B, Litvintseva AP. 2021. Positive Correlation Between Candida auris Skin-Colonization Burden and Environmental Contamination at a Ventilator-Capable Skilled Nursing Facility in Chicago. Clin Infect Dis 73:1142–1148.

10. Vallabhaneni S, Kallen A, Tsay S, Chow N, Welsh R, Kerins J, Kemble SK, Pacilli M, Black SR, Landon E, Ridgway J, Palmore TN, Zelzany A, Adams EH, Quinn M, Chaturvedi S, Greenko J, Fernandez R, Southwick K, Furuya EY, Calfee DP, Hamula C, Patel G, Barrett P, Lafaro P, Berkow EL, Moulton-Meissner H, Noble-Wang J, Fagan RP, Jackson BR, Lockhart SR, Litvintseva AP, Chiller TM. 2017. Investigation of the First Seven Reported Cases of Candida auris, a Globally Emerging Invasive, Multidrug-Resistant Fungus-United States, May 2013-August 2016. Am J Transplant 17:296–299.

11. Omardien S, Teska P. 2024. Skin and hard surface disinfection against Candida auris - What we know today. Front Med (Lausanne) 11:1312929.

12. Borton D. 2024. Candida auris: Emerging fungal pathogen in the US. Nursing 54:23–32.

13. Chen J, Tian S, Han X, Chu Y, Wang Q, Zhou B, Shang H. 2020. Is the superbug fungus really so scary? A systematic review and meta-analysis of global epidemiology and mortality of Candida auris. BMC Infect Dis 20:827.

14. CDC. 2019. Antibiotic resistance threats in the United States, 2019. Atlanta, GA: U.S. Department of Health and Human Services, CDC; 2019.

15. WHO. 2022. WHO Fungal Priority Pathogens List to Guide Research, Development and Public Health Action, 1st ed. World Health Organization, Geneva.

16. Montoya AM. 2024. The Importance of Candida auris in Skin. Current Fungal Infection Reports 18:95–101.

17. Eyre DW, Sheppard AE, Madder H, Moir I, Moroney R, Quan TP, Griffiths D, George S, Butcher L, Morgan M, Newnham R, Sunderland M, Clarke T, Foster D, Hoffman P, Borman AM, Johnson EM, Moore G, Brown CS, Walker AS, Peto TEA, Crook DW, Jeffery KJM. 2018. A Candida auris Outbreak and Its Control in an Intensive Care Setting. N Engl J Med 379:1322–1331.

18. Proctor DM, Dangana T, Sexton DJ, Fukuda C, Yelin RD, Stanley M, Bell PB, Baskaran S, Deming C, Chen Q, Conlan S, Park M, Program NCS, Welsh RM, Vallabhaneni S, Chiller T, Forsberg K, Black SR, Pacilli M, Kong HH, Lin MY, Schoeny ME, Litvintseva AP, Segre JA, Hayden MK. 2021. Integrated genomic, epidemiologic investigation of Candida auris skin colonization in a skilled nursing facility. Nat Med 27:1401–1409.

19. Tharp B, Zheng R, Bryak G, Litvintseva AP, Hayden MK, Chowdhary A, Thangamani S. 2023. Role of Microbiota in the Skin Colonization of Candida auris. mSphere 8:e0062322.

20. Eix EF, Johnson CJ, Wartman KM, Kernien JF, Meudt JJ, Shanmuganayagam D, Gibson ALF, Nett JE. 2022. Ex Vivo Human and Porcine Skin Effectively Model Candida auris Colonization, Differentiating Robust and Poor Fungal Colonizers. J Infect Dis 225:1791–1795.

21. Horton MV, Johnson CJ, Kernien JF, Patel TD, Lam BC, Cheong JZA, Meudt JJ, Shanmuganayagam D, Kalan LR, Nett JE. 2020. Candida auris Forms High-Burden Biofilms in Skin Niche Conditions and on Porcine Skin. mSphere 5.

22. Johnson CJ, Eix EF, Lam BC, Wartman KM, Meudt JJ, Shanmuganayagam D, Nett JE. 2021. Augmenting the Activity of Chlorhexidine for Decolonization of Candida auris from Porcine skin. J Fungi (Basel) 7.

23. Huang X, Hurabielle C, Drummond RA, Bouladoux N, Desai JV, Sim CK, Belkaid Y, Lionakis MS, Segre JA. 2021. Murine model of colonization with fungal pathogen Candida auris to explore skin tropism, host risk factors and therapeutic strategies. Cell Host Microbe 29:210–221 e6.

24. Guolei Zhao JL, Natalia A Veniaminova, Robert Zarnowski, Eliciane Mattos, Chad J Johnson, Derek Quintanilla, Haley Hautau, LeeAnn A. Hold, Bin Xu, Juliet A. E. Anku, Steph S. Steltzer, Kaustav Dasgupta, Darian J. Santana, Ashraf Ibrahim, David Andes, Jeniel E. Nett, Shakti Singh, Adam C. Abraham, Megan L. Killian, J. Michelle Kahlenberg, Sunny Y. Wong, Teresa R. O’Meara. 2025. Candida auris skin colonization is mediated by Als4112 and interactions with host extracellular matrix proteins, bioRxiv.

25. Santana DJ, Anku JAE, Zhao G, Zarnowski R, Johnson CJ, Hautau H, Visser ND, Ibrahim AS, Andes D, Nett JE, Singh S, O’Meara TR. 2023. A Candida auris-specific adhesin, Scf1, governs surface association, colonization, and virulence. Science 381:1461–1467.

26. Shivarathri R, Chauhan M, Datta A, Das D, Karuli A, Aptekmann A, Jenull S, Kuchler K, Thangamani S, Chowdhary A, Desai JV, Chauhan N. 2024. The Candida auris Hog1 MAP kinase is essential for the colonization of murine skin and intradermal persistence. mBio 15:e0274824.

27. Swaney MH, Nelsen A, Sandstrom S, Kalan LR. 2023. Sweat and Sebum Preferences of the Human Skin Microbiota. Microbiol Spectr 11:e0418022.

28. Scharschmidt TC, Fischbach MA. 2013. What Lives On Our Skin: Ecology, Genomics and Therapeutic Opportunities Of the Skin Microbiome. Drug Discov Today Dis Mech 10.

29. Strijbis K, Distel B. 2010. Intracellular acetyl unit transport in fungal carbon metabolism. Eukaryot Cell 9:1809–15.

30. Nev OA, David-Palma M, Heitman J, Brown AJP, Coelho MA. 2024. Fungal pathogens and symbionts: Living off the fat of the land. PLoS Pathog 20:e1012551.

31. Lorenz MC, Bender JA, Fink GR. 2004. Transcriptional response of Candida albicans upon internalization by macrophages. Eukaryot Cell 3:1076–87.

32. Lorenz MC, Fink GR. 2001. The glyoxylate cycle is required for fungal virulence. Nature 412:83–6.

33. Lorenz MC, Fink GR. 2002. Life and death in a macrophage: role of the glyoxylate cycle in virulence. Eukaryot Cell 1:657–62.

34. Miramon P, Pountain AW, Lorenz MC. 2023. Candida auris-macrophage cellular interactions and transcriptional response. Infect Immun 91:e0027423.

35. Ramirez MA, Lorenz MC. 2007. Mutations in alternative carbon utilization pathways in Candida albicans attenuate virulence and confer pleiotropic phenotypes. Eukaryot Cell 6:280–90.

36. Ramirez MA, Lorenz MC. 2009. The transcription factor homolog CTF1 regulates beta-oxidation in Candida albicans. Eukaryot Cell 8:1604–14.

37. Hube B, Stehr F, Bossenz M, Mazur A, Kretschmar M, Schafer W. 2000. Secreted lipases of Candida albicans: cloning, characterisation and expression analysis of a new gene family with at least ten members. Arch Microbiol 174:362–74.

38. Munoz JF, Gade L, Chow NA, Loparev VN, Juieng P, Berkow EL, Farrer RA, Litvintseva AP, Cuomo CA. 2018. Genomic insights into multidrug-resistance, mating and virulence in Candida auris and related emerging species. Nat Commun 9:5346.

39. Naglik JR, Rodgers CA, Shirlaw PJ, Dobbie JL, Fernandes-Naglik LL, Greenspan D, Agabian N, Challacombe SJ. 2003. Differential expression of Candida albicans secreted aspartyl proteinase and phospholipase B genes in humans correlates with active oral and vaginal infections. J Infect Dis 188:469–79.

40. Schwarzmuller T, Ma B, Hiller E, Istel F, Tscherner M, Brunke S, Ames L, Firon A, Green B, Cabral V, Marcet-Houben M, Jacobsen ID, Quintin J, Seider K, Frohner I, Glaser W, Jungwirth H, Bachellier-Bassi S, Chauvel M, Zeidler U, Ferrandon D, Gabaldon T, Hube B, d’Enfert C, Rupp S, Cormack B, Haynes K, Kuchler K. 2014. Systematic phenotyping of a large-scale Candida glabrata deletion collection reveals novel antifungal tolerance genes. PLoS Pathog 10:e1004211.

41. Otzen C, Bardl B, Jacobsen ID, Nett M, Brock M. 2014. Candida albicans utilizes a modified beta-oxidation pathway for the degradation of toxic propionyl-CoA. J Biol Chem 289:8151–69.

42. Findley K, Oh J, Yang J, Conlan S, Deming C, Meyer JA, Schoenfeld D, Nomicos E, Park M, Program NIHISCCS, Kong HH, Segre JA. 2013. Topographic diversity of fungal and bacterial communities in human skin. Nature 498:367–70.

43. Tuor M, LeibundGut-Landmann S. 2023. The skin mycobiome and intermicrobial interactions in the cutaneous niche. Curr Opin Microbiol 76:102381.

44. Ianiri G, LeibundGut-Landmann S, Dawson TL, Jr. 2022. Malassezia: A Commensal, Pathogen, and Mutualist of Human and Animal Skin. Annu Rev Microbiol 76:757–782.

45. Gioti A, Nystedt B, Li W, Xu J, Andersson A, Averette AF, Munch K, Wang X, Kappauf C, Kingsbury JM, Kraak B, Walker LA, Johansson HJ, Holm T, Lehtio J, Stajich JE, Mieczkowski P, Kahmann R, Kennell JC, Cardenas ME, Lundeberg J, Saunders CW, Boekhout T, Dawson TL, Munro CA, de Groot PW, Butler G, Heitman J, Scheynius A. 2013. Genomic insights into the atopic eczema-associated skin commensal yeast Malassezia sympodialis. mBio 4:e00572–12.

46. Wu G, Zhao H, Li C, Rajapakse MP, Wong WC, Xu J, Saunders CW, Reeder NL, Reilman RA, Scheynius A, Sun S, Billmyre BR, Li W, Averette AF, Mieczkowski P, Heitman J, Theelen B, Schroder MS, De Sessions PF, Butler G, Maurer-Stroh S, Boekhout T, Nagarajan N, Dawson TL, Jr. 2015. Genus-Wide Comparative Genomics of Malassezia Delineates Its Phylogeny, Physiology, and Niche Adaptation on Human Skin. PLoS Genet 11:e1005614.

47. Xu J, Saunders CW, Hu P, Grant RA, Boekhout T, Kuramae EE, Kronstad JW, Deangelis YM, Reeder NL, Johnstone KR, Leland M, Fieno AM, Begley WM, Sun Y, Lacey MP, Chaudhary T, Keough T, Chu L, Sears R, Yuan B, Dawson TL, Jr. 2007. Dandruff-associated Malassezia genomes reveal convergent and divergent virulence traits shared with plant and human fungal pathogens. Proc Natl Acad Sci U S A 104:18730–5.

48. Gabriel F, Accoceberry I, Bessoule JJ, Salin B, Lucas-Guerin M, Manon S, Dementhon K, Noel T. 2014. A Fox2-dependent fatty acid ss-oxidation pathway coexists both in peroxisomes and mitochondria of the ascomycete yeast Candida lusitaniae. PLoS One 9:e114531.

49. Kurihara T, Ueda M, Okada H, Kamasawa N, Naito N, Osumi M, Tanaka A. 1992. Beta-oxidation of butyrate, the short-chain-length fatty acid, occurs in peroxisomes in the yeast Candida tropicalis. J Biochem 111:783–7.

50. Chew SY, Chee WJY, Than LTL. 2019. The glyoxylate cycle and alternative carbon metabolism as metabolic adaptation strategies of Candida glabrata: perspectives from Candida albicans and Saccharomyces cerevisiae. J Biomed Sci 26:52.

51. Piekarska K, Mol E, van den Berg M, Hardy G, van den Burg J, van Roermund C, MacCallum D, Odds F, Distel B. 2006. Peroxisomal fatty acid beta-oxidation is not essential for virulence of Candida albicans. Eukaryot Cell 5:1847–56.

52. Strijbis K, van Roermund CW, van den Burg J, van den Berg M, Hardy GP, Wanders RJ, Distel B. 2010. Contributions of carnitine acetyltransferases to intracellular acetyl unit transport in Candida albicans. J Biol Chem 285:24335–46.

53. Jia N, Arthington-Skaggs B, Lee W, Pierson CA, Lees ND, Eckstein J, Barbuch R, Bard M. 2002. Candida albicans sterol C-14 reductase, encoded by the ERG24 gene, as a potential antifungal target site. Antimicrob Agents Chemother 46:947–57.

54. Sprouffske K, Wagner A. 2016. Growthcurver: an R package for obtaining interpretable metrics from microbial growth curves. BMC Bioinformatics 17:172.

55. Dobin A, Davis CA, Schlesinger F, Drenkow J, Zaleski C, Jha S, Batut P, Chaisson M, Gingeras TR. 2013. STAR: ultrafast universal RNA-seq aligner. Bioinformatics 29:15–21.

56. Heinz S, Benner C, Spann N, Bertolino E, Lin YC, Laslo P, Cheng JX, Murre C, Singh H, Glass CK. 2010. Simple combinations of lineage-determining transcription factors prime cis-regulatory elements required for macrophage and B cell identities. Mol Cell 38:576–89.

57. Love MI, Huber W, Anders S. 2014. Moderated estimation of fold change and dispersion for RNA-seq data with DESeq2. Genome Biol 15:550.

58. Xu S, Hu E, Cai Y, Xie Z, Luo X, Zhan L, Tang W, Wang Q, Liu B, Wang R, Xie W, Wu T, Xie L, Yu G. 2024. Using clusterProfiler to characterize multiomics data. Nat Protoc 19:3292–3320.

